# The role of replication-removal spatial correlations and cellular replicative lifespan in corneal epithelium homeostasis

**DOI:** 10.1101/2020.02.26.963199

**Authors:** Lior Strinkovsky, Evgeny Havkin, Ruby Shalom-Feuerstein, Yonatan Savir

## Abstract

Homeostasis in adult tissues relies on the replication dynamics of stem cells, their progenitors and the spatial balance between them. This spatial and kinetic coordination is crucial to the successful maintenance of tissue size and its replenishment with new cells. However, our understanding of the role of cellular replicative lifespan and spatial correlation between cells in shaping tissue integrity is still lacking. We developed a mathematical model for the stochastic spatial dynamics that underlie the rejuvenation of corneal epithelium. Our model takes into account different spatial correlations between cell replication and cell removal. We derive the tradeoffs between replicative lifespan, spatial correlation length, and tissue rejuvenation dynamics. We determine the conditions that allow homeostasis and are consistent with biological timescales, pattern formation, and mutants phenotypes. Our results can be extended to any cellular system in which spatial homeostasis is maintained through cell replication.

## Introduction

In adult tissues, stem cells and their progeny play a crucial role in maintaining homeostasis. Renewal of the tissue is due to progenitor cells that have limited replication capacity (1). The interplay between stem cells and their progenitors with respect to replication, differentiation and cellular hierarchy is not fully understood. For example, two opposing limiting models of stem cell replication have been proposed: A “Hierarchical” model where stem cells are rare slow-dividing cells with longevity similar to the hematopoietic stem cell paradigm (2) and a “Stochastic” model where stem-cells are abundant equipotent cells that divide frequently and their loss is dictated by neutral drift (3–6). Another lingering question is the role of spatial correlation between replication and removal in homeostasis. Some studies assume a long-range correlation between replication and cell removal, that is, as a cell replicates, the removed cell can be tens and even hundreds of cells away (7–10) while experimental studies suggest short-range correlations between replication and removal (11,12).

The cornea acts as a lens that focuses light into the eye, and serves as a barrier that protects the eye against external hazards or injury. Thus, maintaining its integrity and its continuous regeneration is crucial for proper vision in vertebrates (13). It is now predominantly accepted that the regeneration of the corneal epithelium, in homeostasis, is due to limbal epithelial stem cells (LESCs) residing at the circumference of the cornea, the limbus, which separates the cornea from the conjunctiva (Fig. 1A) (14–19). LESCs divide both symmetrically and asymmetrically to yield progenitor cells that have a limited replicative lifespan (RLS) (20). In turn, progenitor cells are proliferating from the limbus to the basal cornea where they can either proliferate or migrate to upper layers (Fig. 1A) (21). The observation that in homeostasis the overall number of cells in the cornea does not fluctuate dramatically led to the ‘XYZ hypothesis’ that states that the proliferation of epithelial cells in the limbus and their migration to the cornea is balanced by cell loss (22). Lineage tracing experiments in living mice revealed a pattern of clonal stripes that propagate from the limbus towards the center of the cornea (Fig. 1B) (23–26). Some hypotheses have been suggested regarding the mechanism of the centripetal migration dynamics in the corneal epithelium homeostasis (27), including population dynamics and electrophysiological or electrochemical cues (28–32). Yet, the underlying mechanism behind these dynamics is not fully understood.

**Figure 1:**
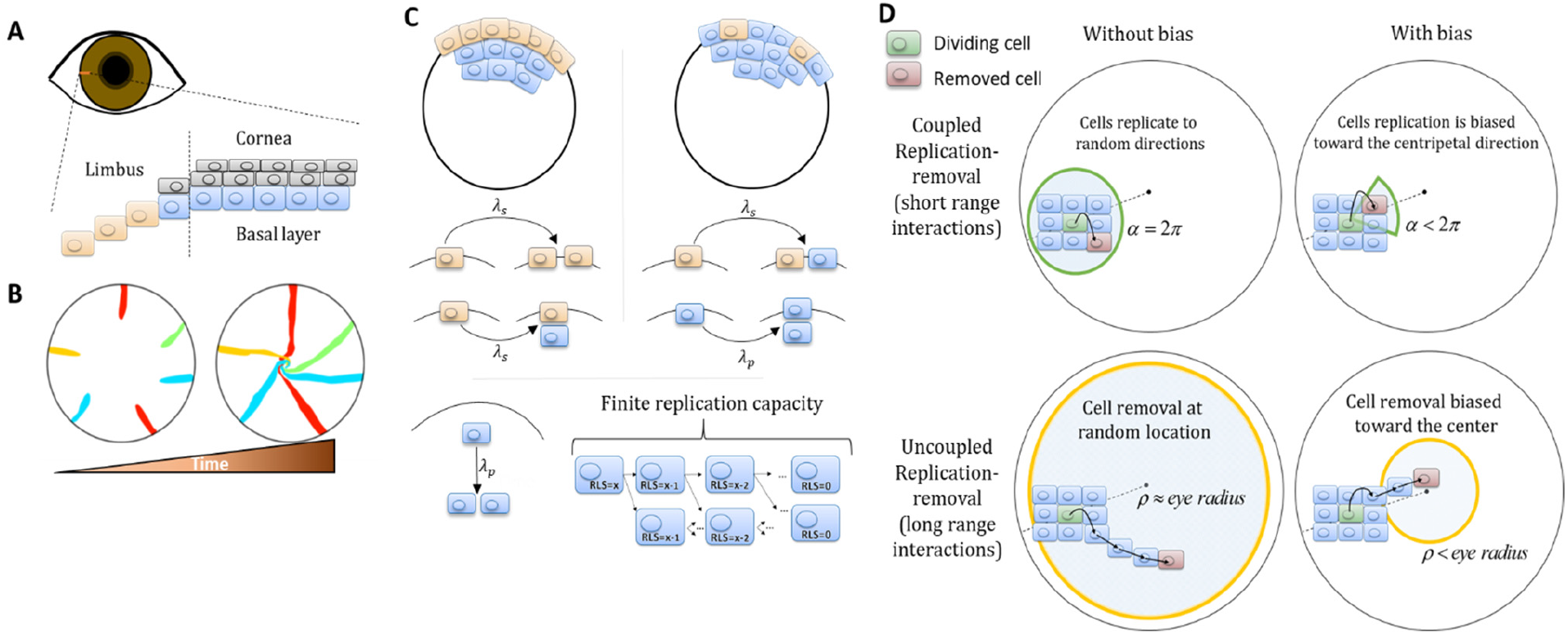
Model setup. **(A)** Maintenance of cells in the cornea, the outer transparent part of the eye, is mainly due to stem cells that reside in the limbus – a niche at the circumference of the cornea. **(B)** Illustration of *in vivo* multi-color lineage tracing experiment. Clonal stripes emerge from the limbus towards the center of the cornea. **(C)** We consider two limiting scenarios for stem cell distribution and dynamics. In the first (left), stem cells are evenly distributed all over the limbus. They can divide symmetrically to maintain the limbus or asymmetrically to provide progenitor cells to the cornea. In the second model (right), stem cells are scarce and divide asymmetrically to give rise to progenitor cells in the limbus that further divide and populate the cornea. In both models, progenitor cells have limited replicative lifespan capacity. When cells exhaust their replicative lifespan, the cells can no longer divide in the basal layer and are replaced by other dividing cells. **(D)** Whether the location of replication and cell removal are correlated (replication and cell removal occur in the same neighborhood) or not plays a crucial role in the cornea rejuvenation dynamics. Accounting also for whether there is a centripetal bias results in four possible classes of models.

Using one-dimensional mathematical models to infer stem cells and progenitor cells dynamics from lineage tracing experiments have been useful in modeling different tissues such as murine epithelial homeostasis of the skin (33), gut (34), human epithelial homeostasis of the epidermis (35) lung (36) prostate (37) and the cornea (38,39). Yet, while these models are insightful, the unique spatial organization in the cornea requires two-dimensional models to encompass overall pattern changes. These types of models require some assumptions on the spatial nature and organization of stem cells and on the spatial interactions between corneal cells. In particular, the role of interaction range between cells, or the spatial correlation between the location of replication and location of cell loss, plays a crucial role in the resulting dynamics (11,12). In addition, another key property of the dynamics is the replicative lifespan (RLS) of corneal cells – the number of times each corneal cell can divide. Approximating the tissue as a kin of elastic network that minimize some energy function (40) was used to suggest that corneal epithelial cells can organize into centripetal patterns in the absence of external cues (41) under the assumption that stem cells are uniformly distributed, RLS of few divisions, and no correlation between replication location and cell loss location.

In this work, we combine a novel mathematical model that allows rapid simulation of the stochastic dynamics of epithelial cells and pattern formation in the cornea together with analytical calculations to consider a broad set of possible physiological scenarios. In particular, we determine the consequences of different assumptions on the replication-removal coupling range and replicative lifespan values. We characterize the tradeoff between renewal time and replicative lifespan and determine the constraints that allow homeostasis and are consistent with the formation of the observed spatial patterns, biological timescales, and mutant dynamics.

## Methods

We developed a lattice-based mathematical model of the cornea’s basal layer, to examine the potential underlying mechanisms and parameters that govern corneal homeostasis, centripetal migration and spatial order patterns as seen in the *in vivo* data (SI section I). We model the cornea as a round assortment of cells in the basal layer of the cornea (Fig. 1C) with a radius *R*. We assume two types of cells: stem cells (*S*) and progenitor cells (*P*) that have different doubling rates *λ*_*s*_ and *λ*_*p*_, respectively. *S* cells reside only in the limbus and can either divide symmetrically or asymmetrically to produce *S* and *P* cells (Fig. 1C). In the case of the ‘Stochastic’ model, the *P* cells can reside only in the cornea while in the ‘Hierarchical’ model they can reside both in the limbus and the cornea, *P*_*L*_ and *P*_*C*_, respectively. *P* cells can divide only symmetrically; if they are in the cornea they can divide in any direction and if they are in the limbus (‘Hierarchical’ model) they can divide only towards the cornea (Fig. 1D). *P* cells are also limited in their maximal amount of horizontal replications-their replicative lifespan, RLS, a parameter which will play a major role in the forthcoming results (Fig. 1C).

Assuming corneal homeostasis, as cells in the cornea and the limbus are dividing, new cells are replacing other cells in the basal layer concurrently. The specific parameters of the replication rate, replicative lifespan, replication direction, and the coupling between the cells affect the spatial dynamics. There are two key properties that play a crucial role in tissue homeostasis dynamics, not only in the cornea. The first is the effective interaction range between replication and removal events. In one limit, the location of the cell that is removed is independent of the replication event. In the other limit, the probability of a cell to be removed will be higher in the area near the new cell (e.g. local pressure). The second property is whether there is an external bias that affects the replication direction or removal location due to, for example, chemical cues, matrix structures or local mechanical perturbation such as blinking. While previous modeling efforts focus only on a particular model, which fits the hypothesis of the study, in this work we systematically account for all of these scenarios and provide the physical limitations, biological implications, and feasibility of each model.

Dynamics are simulated using a stochastic 2D lattice Monte-Carlo approach (SI section I). As a cell divides, the location of the cell that is removed depends on the spatial correlation between these events. In the case that replication and removal are correlated, the location of the removed cell is randomly drawn from a uniform distribution of a circular section that is facing the center of the cornea and has an angle of α and radius of few cells (SI, Fig. 1D). If *α* = 2*π*, there is no centripetal bias, and as *α* is smaller the bias is larger (Fig. 1D). In the case there is no correlation between replication and removal, the removed cell is taken randomly from a circle at the center of the eye with radius *ρ*. As *ρ* is smaller, the centripetal bias increases (Fig. 1D). Once the location of the replicated and removed cells has been determined, the cells in the cornea reorient their location. The probability that a cell will move into the vacant hole depends on its distance from the vector that connects the replicated cell (that causes local stress) and the removed cell (which leaves a vacant space) (SI section I). To track the linage dynamics, each stem cell and its progenies are labeled with the same color.

## Results

### Spatial coupling between cell replication and cell removal

First, we examine the case in which replication and removal processes are spatially coupled in the absence of centripetal bias. We will introduce the centripetal bias in the next section. In this case, the replicated and removed cells are from the same neighborhood which is an order of a few cells and is denoted by *m* (Fig. 1D). Accordingly, the effective step size of clone progression is small, an order of a few cells, hence there is a limit on the maximal distance of cell renewal from the limbus. This distance depends on the RLS. If the RLS is small, then cells will exhaust their replicative capacity before reaching the center of the eye (Fig. 2A, Mov. S1). For example, if the RLS is one (i.e. a *P* cell can divide once and then becomes post-mitotic), in steady-state the renewed cell front will propagate to fill the local interaction neighborhood, *m* (SI section I). As RLS increases, cells can replicate further, and the renewed cell front at steady-state will be closer to the center. The steady-state location of the front is closer to the limbus than what is expected if it propagates an additional one cell towards the center for each replicative lifespan added (the deterministic limit, SI section III). For example, even if the RLS is equal to the radius of the tissue, *R* = 100 in our case, the front still does not reach the center (Fig 2B), because *P* cells lose their proliferative potential already in the periphery.

**Figure 2:**
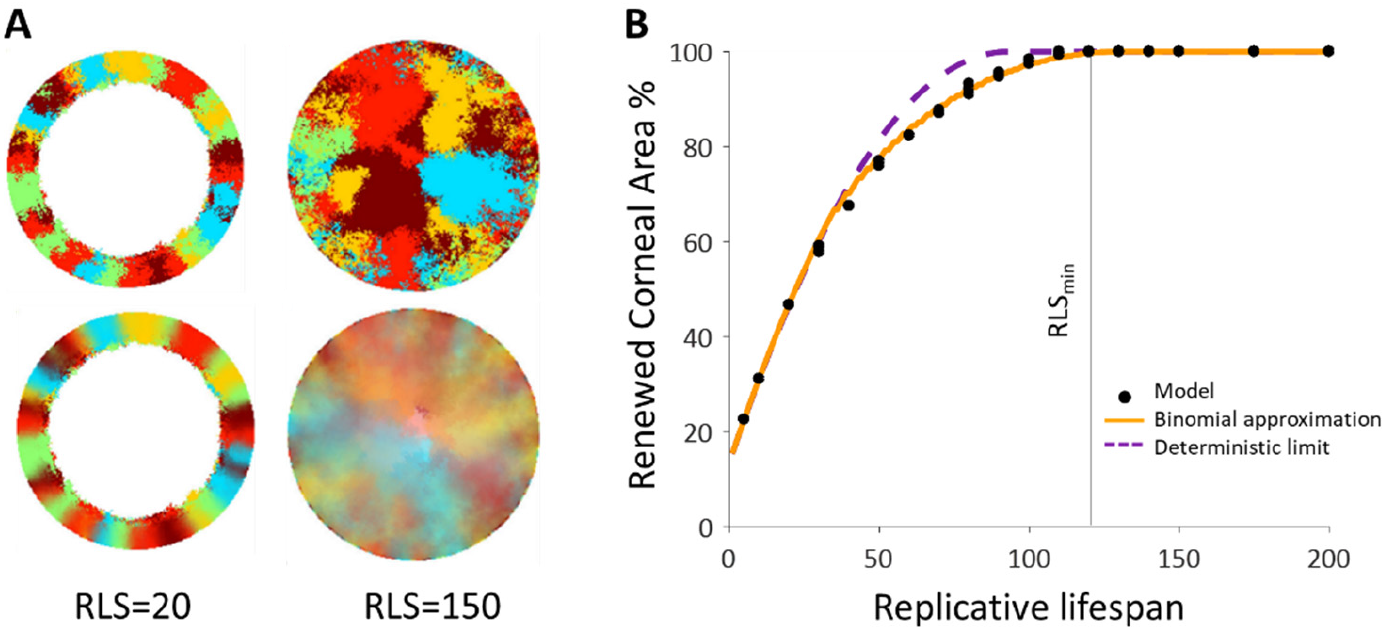
Spatial coupling between cell replication and cell removal in the absence of centripetal bias. **(A)** Steady-state snapshots (upper row) and time-averages over 200 corneal replications (lower row). If the replicative lifespan is below a critical value, the cornea cannot be rejuvenated. For replicative lifespan values that are above the critical value (RLS_min_ ≳ 130), the emerging pattern is that of contiguous patches reminiscent of *in vivo* mutant phenotypes. **(B)** The fraction of the cornea that is renewed increases with replicative lifespan. Three realizations of the dynamics are shown (black dots). The purple line is the theoretical upper limit on the renewed area, and the orange line is the result of a theoretical model that approximates renewal as a one-step process (see text and SI section III for details).

We capture the quantitative details of this phenomenon by considering an effective one-step binomial stochastic process. At each time point, the front can either stay at the same place or move forward with some probability. The probability of moving forward in the case there is no centripetal bias depends on the radial geometry of the front and is estimated to be about 3/8 (SI section III). The results of this model are consistent with the simulation results. In the case there is no centripetal bias the expected minimal RLS, which permits replacement of central corneal cells, RLS_min_, is around 130 replications (Fig. 2B, Mov. S1). For RLSs that are above this critical value, the front can reach the center and a patched pattern is formed. The emerging pattern, in this case, is that of patches (Fig. 2B). While this case does not provide clonal stripes, it does result in contiguous patches that resemble the *in vivo* pattern of mutants that lack certain genes that are thought to play a role in centripetal chemotaxis (42–45).

### Adding bias to the dynamics of coupled replication-removal

We hypothesized that biased cell division orientation towards the center (Fig 1D) would lower RLS_min_ and lead to a pattern that resembles the *in vivo* clonal stripes. To capture the effect of external centripetal bias, (that could be the result of, for example, chemical or mechanical cues), once a cell replicates, the removed cell is drawn randomly from a section that is facing the center of the tissue centered around the replicated cell and has an angle *α* (Fig. 1D). It is convenient to define the centripetal bias in this case to be between zero and one, *bias* = 1-*α*/2π. As the bias increases, a centripetal pattern emerges (Fig. 3A, Mov. S2), and the required minimal RLS for renewal goes down (Fig. 3B). To quantify how much the pattern resembles stripes that elongate from the limbus to the center, we define a clonal unmixing parameter that captures the centripetal stripe mixing (SI section II). When the unmixing parameter is equal to one, the pattern is composed of perfect stripes. As the stripe order is lower, the unmixing parameter approaches zero (SI section II).

**Figure 3:**
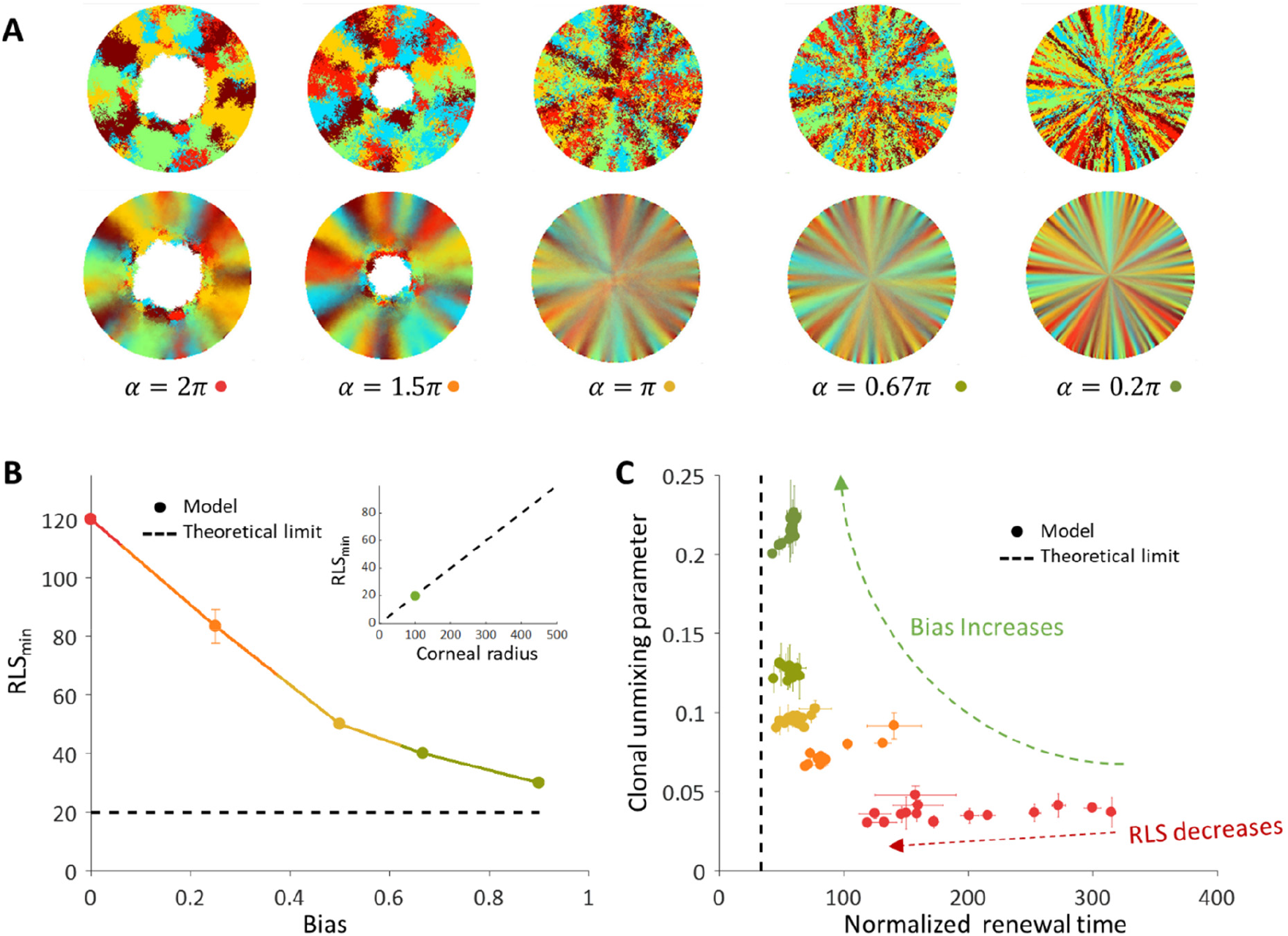
The effect of centripetal bias in the case of spatial coupling between cell replication and cell removal. **(A)** Steady-state snapshots (upper row) and time-averages over 200 replications (lower row) for RLS = 60 and different centripetal bias. **(B)** The minimal RLS required for renewal as a function of centripetal bias. Data points are the mean of three realizations, and error bars (some smaller than the marker size) are the standard deviation. In the case of ideal bias, the minimal theoretical value for RLS_min_ (dotted horizontal line) is ≈ *R/m* (where *R* is the radius of the cornea, and *m* is the local replication-removal neighborhood). (B-inset) The lower limit on RLS_min_ as a function of the corneal radius when *m* =5. The green dot marks the corneal radius used for the simulations, *R* = 100. **(C)** The effect of centripetal bias (Different colors, match the colors on panel 3B) on the normalized renewal time and clonal unmixing. As the bias increases, the corneal renewal time is shorter, and the pattern is more ordered. In the case of ideal bias, the theoretical lower limit on renewal time (vertical dotted line) is 2*R*/(*m*+1).

When there is no bias, the minimal RLS is of the order of the radius of the tissue. In the case of ideal bias, the minimal replicative lifespan required for full renewal, where *m* is much smaller than *R*, is *R*/*m* (SI section III). In our case *m* is five cells which give a minimal RLS of about 20 replications (Fig, 3B, 3B-inlet). This result is interesting in light of previous literature that attributed very limited replication capacity (RLS around 3-4) to short-lived *P* cells (41,46,47). It imposes a minimal lifespan for progenitor cells that scales as the radius of the tissue divided by the radius of the local neighborhood in which cells interact.

The replicative lifespan also plays a role in determining the normalized renewal time that is defined as the time it takes for all cells in the cornea to be replaced divided by the doubling time of corneal cells. As the RLS increases, the renewal time decreases (Fig. 3C). As the centripetal bias is larger the renewal time is faster. In the case of ideal bias, the limit on the renewal time dynamics can be captured as a one-step process with an average step size that depends on the interaction length, *m* (SI). The limit on the renewal time is given by 2*R*/(*m*+1) (Fig. 3C). In our case, where the radius is about 100 cells, it results in a renewal time of ~35 replications, which amount to ~100 days assuming a 3-day cell cycle (46,48).

### Spatial uncoupling of replication-removal

Spatial coupling between cell replication and cell removal has been demonstrated in skin cells, yet another possibility is the case where replication and removal are not tightly correlated in space. In this case, the replication and removal do not have to be in the same neighborhood (Fig 1D). First, we consider the case where there is no inherent centripetal bias, that is cell removal and replication events in the cornea are not biased toward the center. It was previously suggested that centripetal patterns can be formed even in the absence of centripetal bias (7). We show here that this phenomenon is limited to a particular set of RLS and interaction lengths that are associated with slow corneal replenishment time and high post-mitotic rate.

The emergent pattern, in this case, is inherently different. For low RLS values, a centripetal pattern is formed near the limbus edge. However, the unmixing is diminishing as the stripes are moving towards the center (Fig. 4A, 4B, Mov. S3), similar to previous reports(41). The unmixing is decaying towards the center of the tissue and the pattern becomes mixed akin of a ‘salt and pepper’ noise. In this case, the disordered pattern does not form spatial neighborhoods as the coupled interaction case.

**Figure 4:**
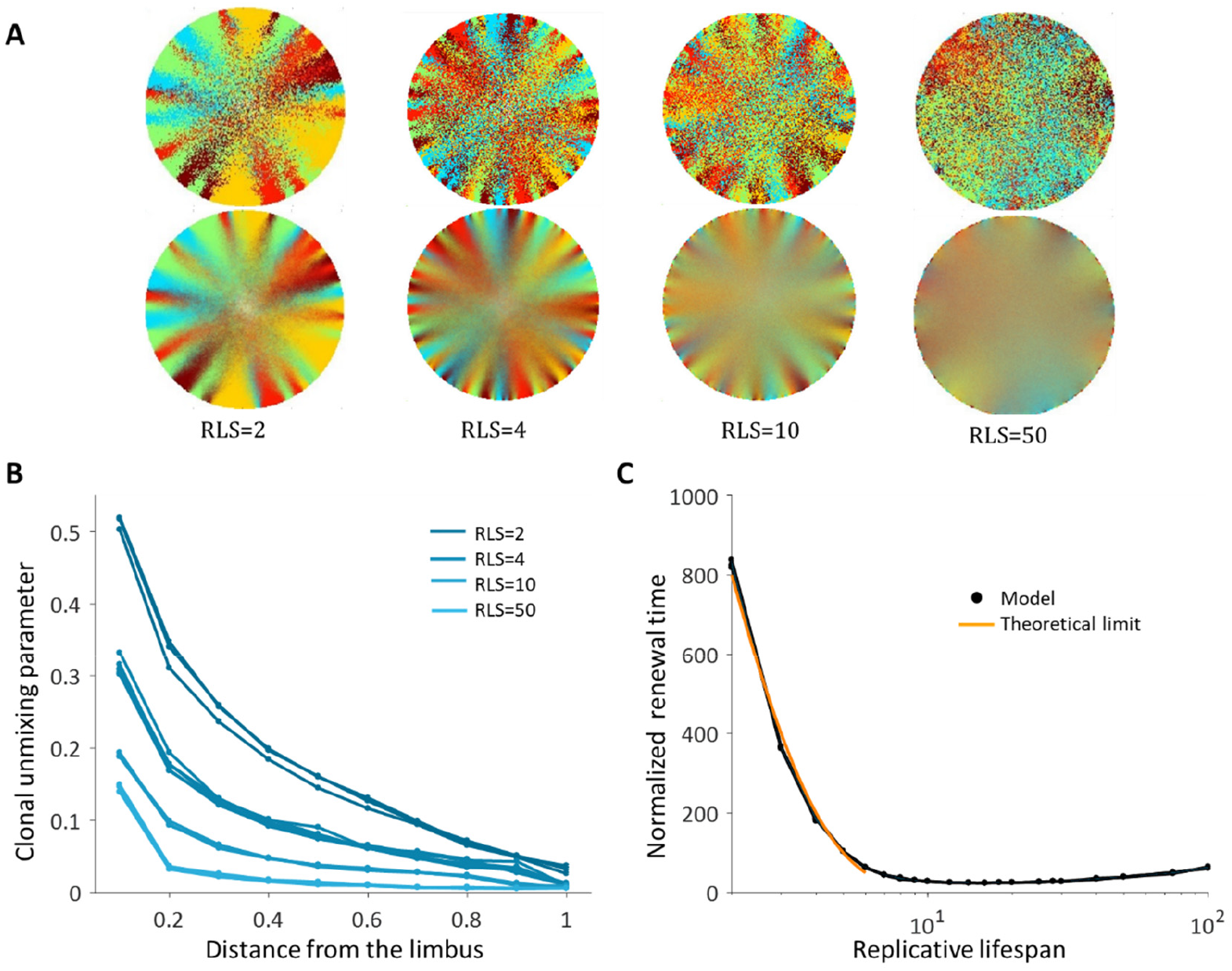
Spatial uncoupling between cell replication and cell removal. **(A)** Steady-state snapshots (upper row) and time-averages over 200 corneal replications (lower row) for different RLS values. **(B)** The clonal unmixing at different distances from the limbus. Ordered centripetal pattern emerges mainly at the periphery of the tissue and only for low values of replicative lifespan (blue-lines, three realizations for each RLS value). **(C)** The renewal time decreases exponentially with RLS for values that are lower than log_2_(*R*) ≈ 7 (black line, three realizations are shown). The orange line is the theoretical limit when considering renewal as a one-step stochastic process with radial boundary (see text and SI section IV).

The emergence of centripetal pattern occurs only for small RLS values. As the replicative lifespan is larger than a few replications, the emergent unmixing becomes more and more limited to the edges of the tissue. This is consistent with previous studies that assumed RLS of only a few replications. For RLS that is larger than ten, unmixed order formation is very limited. In contrast, small RLS values result in a high post-mitotic rate (Figure 7B). Interestingly, while only very low RLS values allowed some organized stripe pattern, in these low RLS values, the vast majority of the cornea was occupied by post-mitotic *P* cells that continuously underwent centripetal movement as a consequence of divisions of limbal *S* and corneal peripheral *P* cells. Thus, this regime suggests no cell proliferation at the center of the cornea.

Another ramification of short replicative lifespan is the number of replications needed to get to the center of the tissue (replication times) that also depends greatly on the RLS. When the capacity of the cells to replicate is low, RLS of only a few replications, the time that is required to get to the center in units of replication times is in the order of hundreds (Fig. 4C). Thus, there is an inherent tradeoff between the renewal time and the centripetal order.

It is insightful to estimate the limits on the maximal and minimal renewal times in this case. For the maximal time: In case the RLS is zero, that is only the stem cells can replicate, the fastest renewal time is when there is a perfect bias towards the center. In this case, the renewal time is determined by the radius of the tissue, the asymmetric replication rate of the stem cells and the geometrical difference between the number of cells that are replicating to the number of cells that are pushed forward (SI section III). In terms of the number of corneal cells replications, *t*_*with bias*_ = (*λ*_*p*_/(2*λ*_*s*_ ⋅ *a*))(*R* + 1), which is about 600 corneal replication times in our case. In the case there is no bias at all, the probability to move towards the center is not one and depends on the location of the front (SI section IV). In this case, the renewal time is larger, *t*_*no bias*_ = (*λ*_*p*_/(*λ*_*s*_ ⋅ *a*))⋅*H*_*R*_ ⋅ *R*, where *H*_*R*_ is the R^th^ harmonic number. In the case the radius is 100 cells, the replenish time increase by a factor of ~10 to ~6000 corneal replication times. As the replicative lifespan increases, additional corneal cells contribute to the propagation of the front. Thus, for small RLS the replenishing time is expected to decrease by the factor 2^*RLS*^, the number of cells that contribute to pushing front. This is consistent with the results of the simulated dynamics (Fig. 4C). For the minimal time: In case of ideal bias and high RLS, in each corneal replication time the traced stripes double their length. Thus, the minimal time needed to renew the whole cornea is log_2_(*R*) which in our case is around 7.

### Adding centripetal bias to the uncoupled model

In the previous section, we show that if cell replication and removal are not coupled in space, the emergent clonal stripe pattern is limited to the periphery and also requires very low RLS values that lead to post-mitosis of most P cells at the corneal periphery. For these values of low RLS, the renewal time is slow and requires hundreds of corneal replications. To study the effect of centripetal bias in this case, we keep the assumption that the location of replication and removal are independent, but the location of the removed cell is from a circle that is centered at the center and has a smaller radius (Fig. 1D). This could result from, for example, localized high pressure in the center of the eye, or from blinking that affects more the cells in the center of the eye. We note that the area with high probability for cell removal does not have to be a circle, for example, blinking can cause a horizontal, elliptic, area of high removal probability (49–51). Here, we assume a circle to capture the qualitative tradeoffs of increasing the bias on the dynamics.

As the bias increases, the overall clonal unmixing increases (Fig. 5A, 5B, Mov. S4). Yet, the overall trend of mixing order in the central region from which cells are removed is bias independent. RLS smaller than ~5 provide high unmixing but results in slow renewal dynamics (Fig 5B), and still high post-mitotic rate (Fig. 6B). It is insightful to consider two types of timescales: one is the time it takes for stripes to reach the center of the tissue, that is an important experimental observable, and the second is the overall renewal time which is the time that takes to fill the entire cornea (Fig. 5B). These two timescales exhibit different dependencies on the replicative lifespan. As the RLS is larger, the time it takes a clone stripe to reach the center is increasing exponentially with RLS. For large values of RLS, as the bias is larger, the velocity of stripe progression is larger. The limit on stripe speed can be estimated In the case of ideal bias, that is the removed cells are from the center of the tissue, and high replicative capacity, the minimal number of replication needed for a stripe to reach the center is given by log_2_(*R*). (SI section IV), where *R* is the radius of the tissue, which is around 7 corneal replication times in the case *R* = 100. The renewal time of the entire tissue dynamics exhibits a non-monotonic behavior. For high RLS values, the centripetal motion slows down the motion in the direction which is orthogonal to the centripetal direction and thus slows down the overall renewal dynamics. The interplay between the renewal time and unmixing is shown in Fig. 5C.

**Figure 5.**
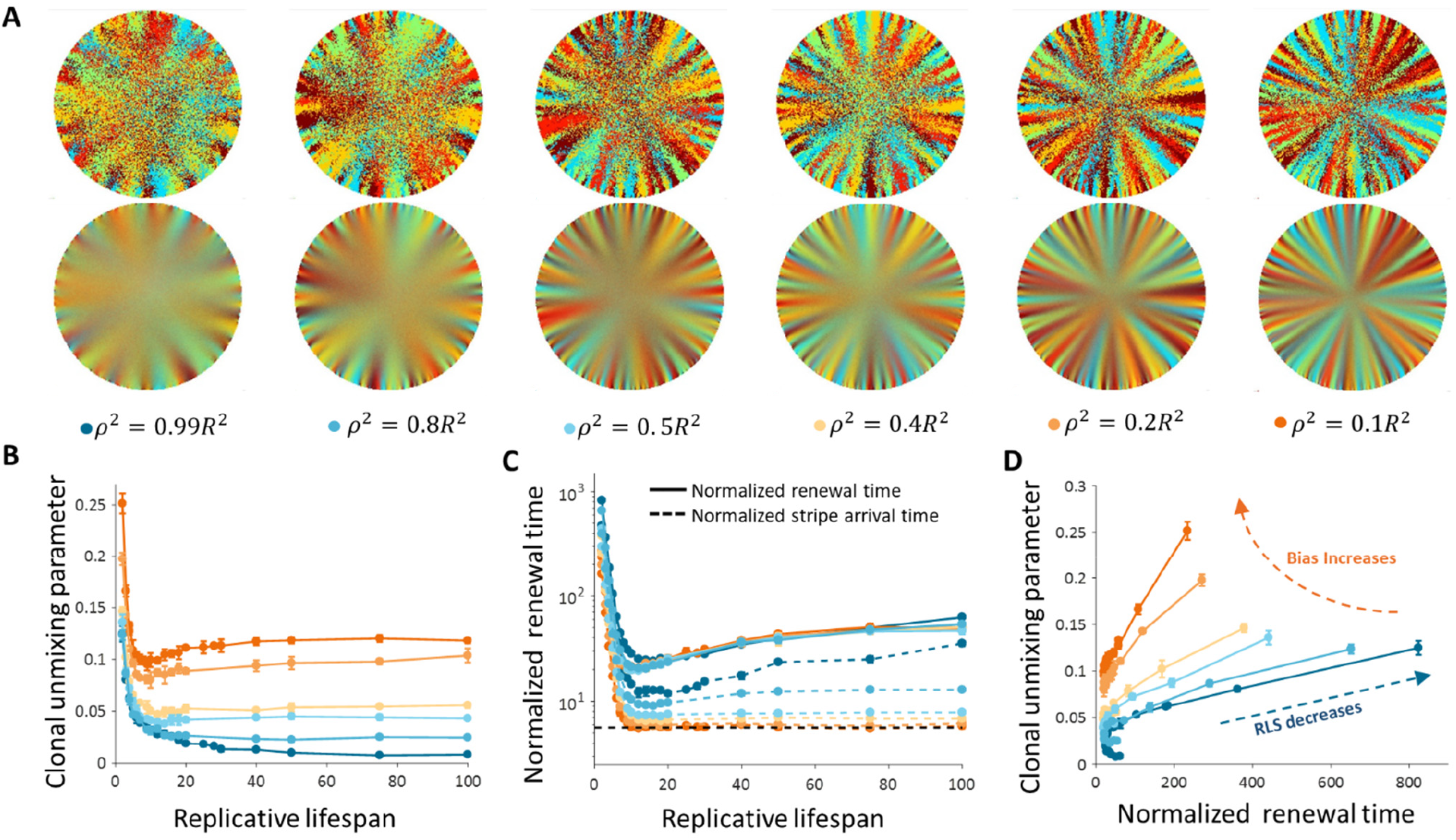
Dynamics and patterns in the case where replication and removal are uncoupled. **(A)** Steadystate snapshots (upper row) and time-averages (lower row) over 200 corneal replications for different values of *ρ*, the radius of the area from which cells can be removed, for RLS = 10. **(B)** The dependence of centripetal unmixing, halfway to the center, on replicative lifespan. Colors denote the centripetal bias and are the same as in 5A. Data points are means of three realizations, and error bars are the standard deviation. For all values of RLS, the order decreases as RLS increases. **(C)** Normalized renewal time decreases as RLS increases (solid lines). The black dotted line is the theoretical limit in the case of ideal bias and high RLS. Data points are means of three realizations, and error bars are standard deviation. **(E)** The interplay between unmixing and renewal time for different bias (different colors) and different RLS values ranging from 2 to 100. Data points are means of three realizations, and error bars are standard deviation.

**Figure 6.**
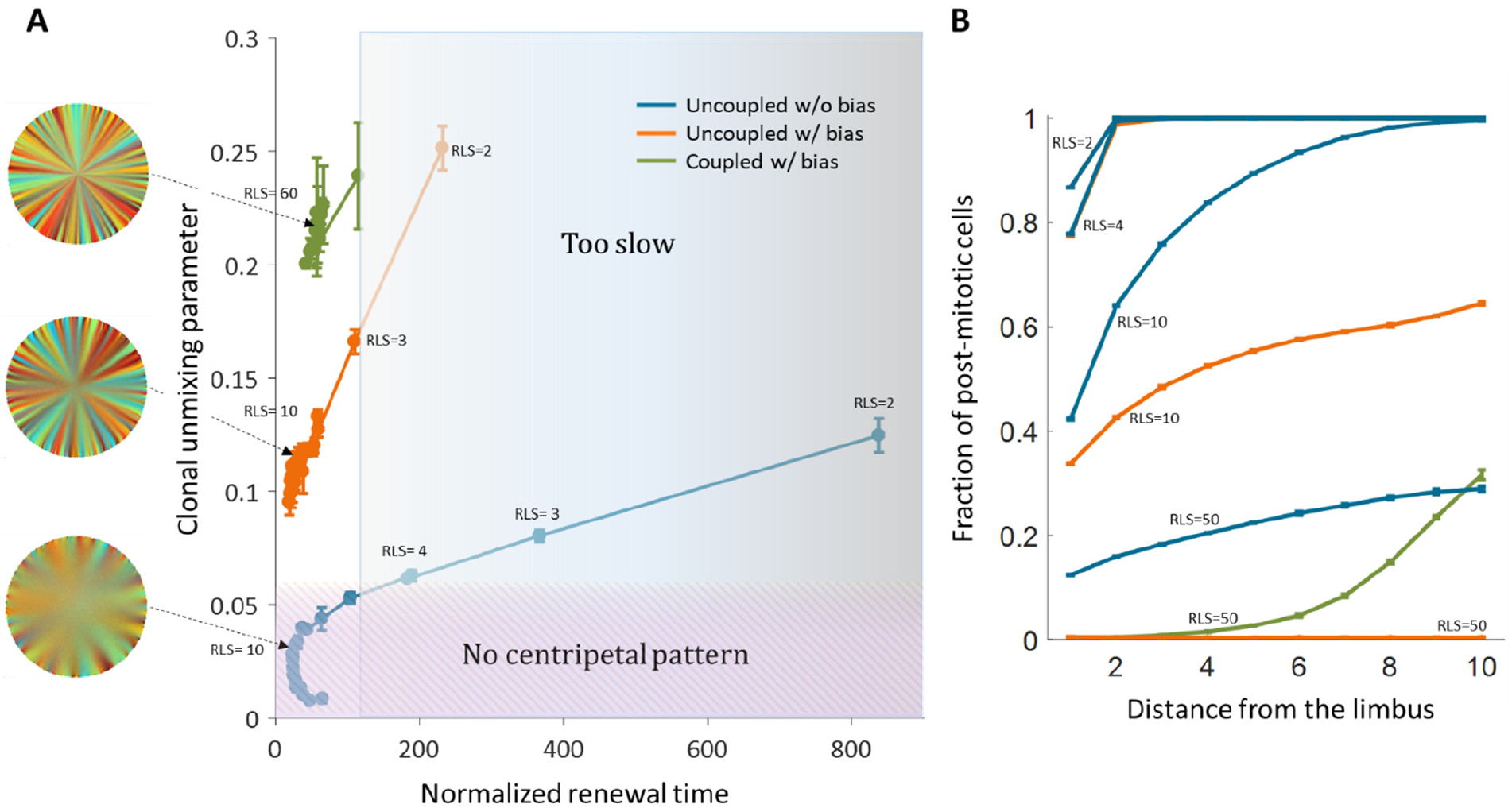
Limits and constraints on the different model classes. **(A)** The unmixing parameter as a function of the normalized renewal time for different values of replication lifespan, ranging from 2 to 100. The shaded areas are regions that are not consistent with experimental observations of centripetal unmixing and renewal time. **(B)** The hallmark of low RLS models is a high post-mitotic fraction regardless of bias.

### Limits on the different models

We have considered three main properties that affect the dynamics and pattern formation in the cornea: spatial correlation between replication and removal, inherent centripetal bias, and replicative lifespan of cells in the cornea. For different models, there were different constraints that determine their biological feasibility. Under conditions of spatial coupling between cell replication and cell loss, external cue that imposes a bias of centripetal movement is required for the emergence of a centripetal pattern. Another major constraint is a requirement for a minimal value of RLS that allows renewal. This value depends on the replication bias and cannot be shorter than a lifespan of ~20 replications. One the condition for the minimal RLS is met, dynamics are feasible, and the emerged unmixing is higher than the unmixing of long-range interactions. In the case of uncoupled interactions without bias centripetal unmixing formation is feasible under constraints. Centripetal clonal unmixing emerges but only on the periphery of the tissue and only for small RLS values that are below~10. However, for these RLS values, the renewal dynamics and stripe velocity, of a few hundred corneal replications, is much slower than observed in experiments which is an upper limit is ~100 replications (Fig. 6A). Adding bias to these types of interactions allows formation in feasible time scales and the emergent of a centripetal pattern for slightly lower RLS values (Fig. 6A).

Another experimental observation is the fact that in some mutants that abolish centripetal bias, the resulting pattern is that of contiguous patches. Models of uncoupled interaction cannot provide such a pattern while coupled can. In the case of uncoupled interactions without bias, the resulting pattern would be akin to ‘salt and pepper’ mixed pattern rather that of patches. Thus, only the coupled model can explain contagious patches in mutants with the constraint of RLS >20. Another consequence of the low replicative lifespan is the distribution of post-mitotic cells. Fig. 6A shows the fraction of post-mitotic cells as a function of the distance from the limbus. Models that require low replicative lifespan, such as uncoupled replication-removal without bias, results in a large portion of the cornea that is post-mitotic.

### The effect of stem cells distribution and dynamics

There are two limiting common hypotheses for the properties of stem cells, and in particular limbal stem cells, in their niche. In the previous section, we investigated the case where stem cells populate all the limbal cells and can replace each other as they replicate. In this section, we also consider the case in which stem cells are rare cells (~10%) (52) which divide asymmetrically to limbal progenitor cells that in turn, divide into corneal cells. In this case, the stem cells cannot be replaced by limbal progenitor cells. The interplay between the emergent pattern and the dependence on replicative lifespan and replication-removal interaction length is similar to the case of uniform progenitor cells (Fig. 7A, 7B).

**Figure 7.**
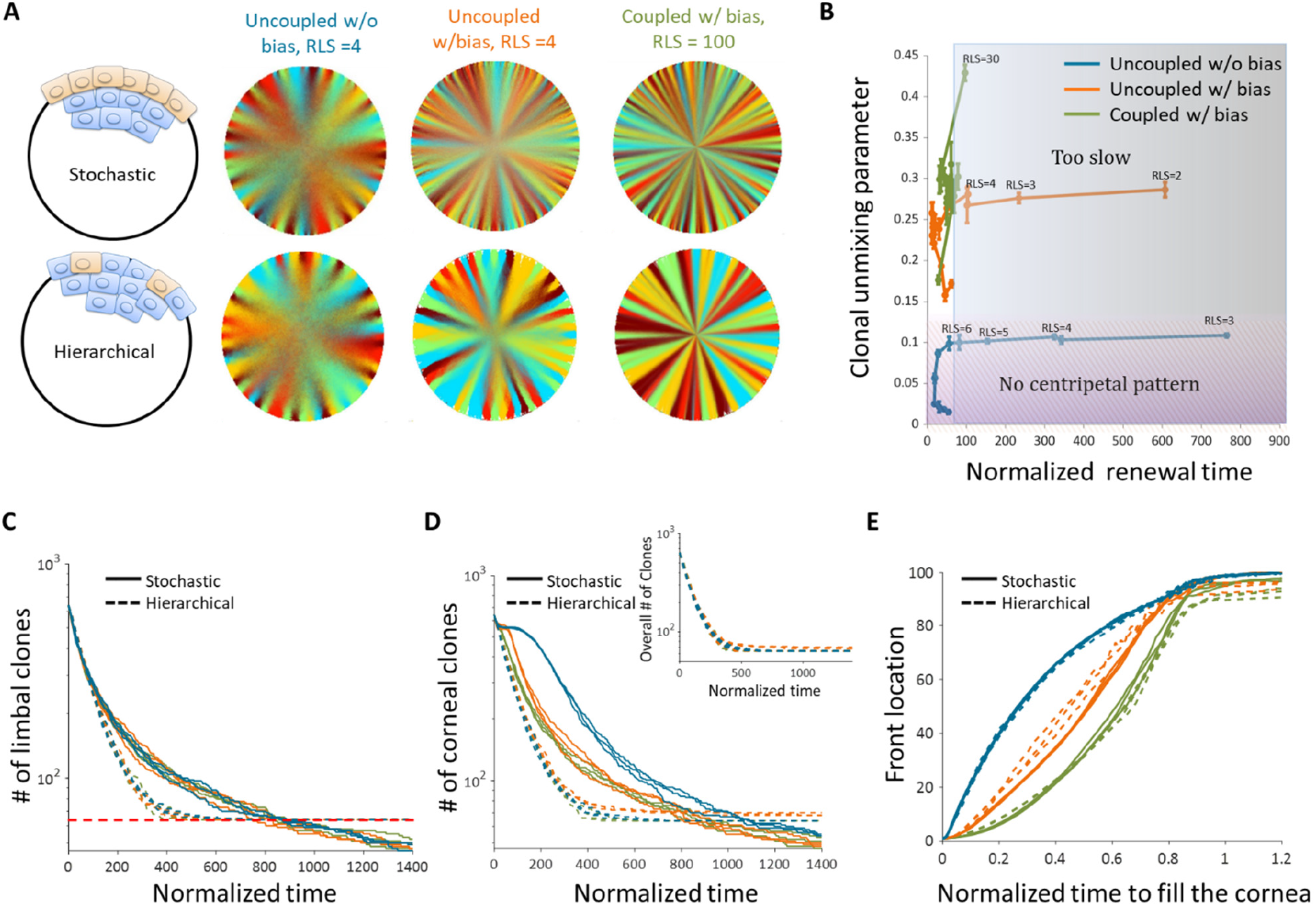
The effect of stem cell dynamics and distribution. **(A)** Steady-state snapshots (upper row) and time-averages (lower row) over 200 corneal replications for the stochastic and hierarchical models. The colors denote the different cases (blue, orange, and green) and are the same for all the panels. **(B)** The interplay between clonal unmixing and renewal time for different bias and different RLS values in the case of the hierarchical model. The tradeoffs are similar to those of the stochastic model (Fig. 6A). **(C)** The number of limbal clones as a function of time in the case of the stochastic model (solid lines) and the hierarchical model (dotted lines). While the number of limbal clones in the stochastic case diminishes with time, the number of limbal clones in the hierarchical case approaches the number of limbal stem cells (red horizontal dotted line). In both models, the spatial coupling does not affect the dynamics of clone number. **(D)** The number of corneal clones as a function of time in the case of the stochastic model (solid lines) and the hierarchical model (dotted lines). In the hierarchical case, the dynamics of corneal clone number and limbal clone number are similar (inset). In the stochastic case, the spatial coupling affects the decay rate of the number of clones. **(E)** The renewed cells’ front location as a function scaled time. Time was scaled such that 1 is the time to replenish the cornea. The stripe propagation velocity depends on the spatial coupling but is less sensitive to the stem cells distribution.

To highlight the differences between these two models, it is insightful to examine the number of clones as a function of time both in the limbus and in the cornea. In the case of the hierarchal model, both the number of clones in the limbus and in the cornea goes down in a similar fashion until it is equal to the number of stem cells. (Fig. 7C). These dynamics are invariant to the interaction length and degree of centripetal of bias. In the case of the stochastic model, the dynamics of the number of clones in the limbus and in the cornea is different. In the limbus, there is a monotonic decline in the number of clones. This decline is invariant to the interaction length due to the continuous competition in the limbus. The dynamics of the number of clones in the cornea has a bi-phasic shape that does depend on the interaction length (Fig. 7D). The first part of the bi-phasic dynamics is a decline that is similar to the limbus decline and is due to stem cell competition. A plateau follows this decline as clones are propagating towards the center. The second decline is due to the clonal competition when all the tissue is ‘colored’. Another experimental measurement is the location of the clone front. The normalized front propagation velocity is invariant to the stem cell dynamics (whether it is the hierarchal or stochastic model) and depends mainly on the interaction length and bias (Fig. 7E).

## Discussion

The dynamics of corneal stem cells and their progenitors play a key role in maintaining homeostasis in adult tissues. As the total number of cells in homeostasis remains constant, the main facilitators of cell location and tissue rejuvenation, when the tissue is intact, are cell division and removal. Thus, the spatial correlation between the locations of the replicated cell and the removed cell determines the rejuvenation speed and clonal pattern of the tissue. Another feature that is critical in replication-removal dynamics is the replicative lifespan of the progenitor cells. Here, we use a mathematical model together with analytical benchmarks to derive the tradeoffs and constraints of varying replication-removal correlation length and replicative lifespan and characterize the conditions that are consistent with experimental measurements. Identification of the conditions governing corneal cell dynamics will facilitate new approaches to limbal stem cell deficiency treatments and translate to other cellular systems that are dependent on spatial cell arrangement and division.

Spatial coupling of replication and removal dramatically influences the parameters that are needed for tissue renewal in physiological time scales. Recent studies suggested that replication and removal events in homeostasis happen in close proximity of few cells (11,12). The main consequence of this type of ‘short-range’ interaction is that they set a minimal replicative lifespan (Fig. 3B). The limit for replicative lifespan is the ratio between the radius of the cornea and the radius of the local neighborhood in which replication and removal occur. For example, in the case of a cornea with a radius of ~100 cells and a local interaction neighborhood with a radius of ~5 cells, the minimal replicative lifespan should be at least 20 replications. This limit is in the case of high bias, that is the cells are replicating towards the center of the tissue. In practice, one should expect a higher limit (Fig. 3B). This suggests that if cell replication and cell removal are indeed spatially correlated, the replicative lifespan of progenitor cells should be much higher than traditional values which are an order of only a few replications. Another interesting consequence of these results is that increasing the tissue size (e.g. human and large mammals cornea and large organs) requires increasing the replicative lifespan of progenitor cells or increasing the local replication-removal interaction length. Cancer, aging or other hyperplastic conditions (e.g. psoriasis) are extreme examples of potentially extensive changes in replicative lifespan that may lead to failure in maintaining tissue renewal and proper tissue size, leading to a burden on stem cells, failure to maintain homeostasis, and/or regenerate the tissue under stress.

We also characterized the dynamics in the limit in which replication and removal events are not spatially correlated. This type of dynamics was suggested in the context of the cornea with a replicative lifespan of fewer than about five replications (7). Our results show that while rejuvenation of the entire cornea is possible for short replicative lifespan, the rejuvenation time without external bias is much slower than physiological estimations (Fig. 6A). Thus, self-organizing stripe formation in homeostasis without external cues, while possible, is very limited and results in a rejuvenation time of hundreds of replications, that is, hundreds of days assuming corneal replication time of a few days. Another hallmark of a model that has a short replicative lifespan is a cornea in which most of the cells are post-mitotic (Fig. 6B).

Figure 8 summarizes the predictions of each model and its consistency with experimental data. We focused on four main experimental attributes: 1) Tissue renewal: whether the model allows complete rejuvenation of all cells in homeostasis. 2) Feasible dynamics: Whether the speed of clonal spread is physiological. 3) Centripetal pattern: Whether the model allows the formation of centripetal clonal stripes. 4) Contiguous patches: Whether the model allows the formation of contiguous clonal patches that is reminiscent of VNGL/PAX6 mutants. The model that seems to account for all features is that of coupled replication-removal dynamics (‘short-range interactions’) with centripetal bias and a replicative lifespan that is at least ~20 replications. One of the main predictions of such a model is that cells should proliferate not only near the limbus but also closer to the center of the cornea.

**Figure 8:**
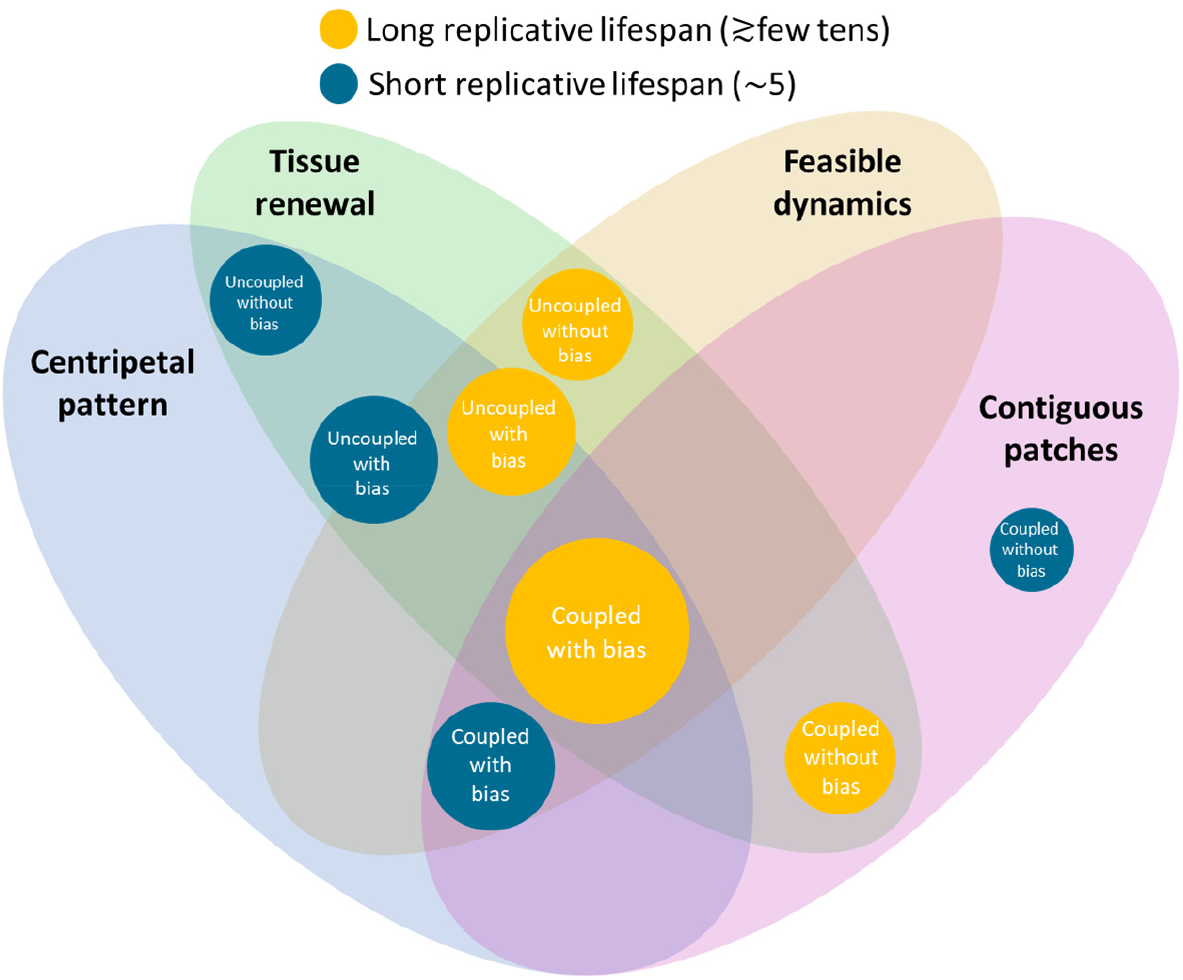
Constraints on the different models and their biological feasibility. Each circle represents a model with different spatial correlations between cell replication and cell removal, high or low replicative lifespan, and whether there is a centripetal bias or not. The colors represent whether the replicative lifespan is short or long (blue and orange, respectively). The radius of each circle is proportional to the number of properties each model is consistent with. In the case of the cornea, the only model that can provide all four requirements (centripetal pattern, full tissue renewal, feasible time scales, and contiguous patches observed in mutants) is coupled replication-removal with bias and long replicative lifespan.

Our results regarding the interplay between replication-removal interaction length and replicative lifespan do not depend on whether stem cell dynamics and distribution follow the ‘Hierarchical’ or ‘Stochastic’ model (Fig. 7B). As expected, our results show that the number of clones overtime on the limbus and cornea together could distinguish between the two. While in the ‘Stochastic’ model, the number of clones in the cornea has a certain plateau and delay between the cornea and the limbus while in the ‘Hierarchical’ model there is no difference in the dynamics of the number of clones in the limbus and in the cornea (figure 7C-D).

While stem cells, that are considered as long-lived cells that can self-renew, are at the focus of regenerative medicine, progenitor cells, that are viewed as short-lived cells with a very limited replication potential, are often overlooked. Our work highlights the crucial role of replicative lifespan of progenitor cells in shaping rejuvenation dynamics in homeostasis. Our conclusions regarding the interplay between replication-removal locality and replicative lifespan are relevant for any tissue in which conditions do not permit significant cell motility and thus spatial homeostasis is maintained through cell replication.

