## Supplementary information for "The role of replication-removal spatial correlations and cellular replicative lifespan in corneal epithelium homeostasis"

#### I. Stochastic 2D lattice Model

**Model setup**. We modeled the corneal epithelium as a circle on a square 2D lattice, where each pixel is a cell. The limbus is a one-cell (that is, a one-pixel wide ring ) perimeter in the circumference of the cornea. Stem cells (*S*) reside in the limbal region in the circumference of the cornea. They are also characterized by immense replicative capacity rendering them 'immortalized' for the sake of the model. Progenitor cells (*P*) that reside in the cornea are limited in their replicative lifespan (RLS): after a defined number of divisions, they cease to divide on the corneal plane.

In the case of the 'Stochastic' model (Fig. 1A), the entire limbus is composed out of *S* cells that divide in the limbus with a rate $\lambda_s$. The cells can divide asymmetrically with probability $p_a$ and give rise to a *P* cell that resides in the cornea or divides symmetrically with a probability of (1- $p_a$), Both *S* cells remain in the limbus. In the cornea, *P* cells replicate with a rate $\lambda_p$. The cells that originate from the stem cells, have RLS replications (Fig. 1B). That is, the maximal number of cells from each *P* cell is $2^{RLS}$.

In the case of the 'Hierarchical' model (Fig. 1A), only a small fraction of cells in the limbus, $f_S$, is *S* cells. These cells are distributed uniformly in the limbus. The *S* cells can divide asymmetrically with a rate $\lambda_S$. The resulting progenitor cells, $P_L$, remain in the limbus and divides with a rate $\lambda_p$ to give rise to a limbal and corneal *P* cells, $P_L$ and $P_C$, respectively. Both $P_L$ and $P_C$ have finite RLS.

We used *R*=100 cells to match the *in vivo* mouse data (Table S1). A distance of 50 pixels was taken from the central pixel to define the pixels of the boundary (that is the horizontal and vertical axis contains 101 pixels). The one-pixel wide ring in the circumference is marked as the limbus. The cornea is maintained in homeostasis. Thus, cell number is constant and for each cell division, there is concurrent cell desquamation. The dynamics of spatial rearrangement of cells in the grid are based on a "pushing" mechanism where the dividing cell creates "pressure" in the direction of the exiting cell and push each cell in the way between them towards the removed cell location (see details below).

**Simulation steps.** We used a 2D Monte-Carlo (MC) simulation. In each MC step of the simulation, a replicating cell and a removed cell are selected as described below. *P* cells exceeding their RLS cannot be selected for division. The probability to select a cell for replication depends on the replication rates according to the Gillespie algorithm (1–3).

The selection of the removed cell depends on the model class: whether the replication-removal are coupled or uncoupled (short-range or long-range interaction), and on the magnitude of the centripetal bias (see details below).

1. If the replicated cell is a corneal cell, that is a *P* cell (in the stochastic case) or a $P_C$ cell (in the hierarchical case) the removed cell will be in the cornea as well. The location of the removed cell will be:
    a. Coupled replication-removal: Randomly selected from a circular sector with a radius $m$ and an angle $\alpha$. The center of the circle is the replicated cell, and the sector is oriented towards the center of the cornea (Fig. 1D).
    b. Uncoupled replication-removal: Randomly selected from a circle around the center of the cornea and has a radius $\rho$ (Fig. 1D).
    c. After a dividing cell and an exiting cell are selected**:**
        i. The dividing cell generates a *P* cell that is in a random direction from the dividing cell and in distance of 0.5 cell grid distance off-grid.
        ii. The algorithm finds the path of cell replacement closest to a straight line that connects the dividing cell and the ‘hole’ that is left on the removed cell location.
        iii. The path is based on 8 directional movements on the square lattice. Excluding directions that make the path cross the limbus of the cornea or exit the boundaries of the cornea.
        iv. On the generated path, the cell closest to the hole moves into it creating a new ‘hole’. The next cell on the path that is closest to the hole moves into it and so on until the new progenitor cell which was off-grid replaces the last cell in the path.

2. If the replicated cell is a limbal cell (*S* in the stochastic case; *S* or $P_L$ in the hierarchical case):
    a. In the stochastic case:
        i. *S* cells divide symmetrically with probability $p_a$. In this case, the *S* offspring remains in the location of the parental *S* cell in the limbus and the *P* offspring becomes a corneal cell. The removed cell will be from the cornea and the location of the removed cell and the rearrangement of cells location is the same as in 1.

ii. $S$ cell divides symmetrically with probability ($1 - p_a$). In this case, the removed cell is chosen randomly from the limbus. The rearrangement of the limbal cells follows the same steps as in 1c, with a path that confined to the limbus.

b. In the hierarchical case:

i. $S$ cells divide asymmetrically. The $S$ offspring remains in the parental $S$ cell in the limbus and the $P_L$ offspring remains a limbal cell. The removed cell is chosen randomly from the $P_L$ cells that are between the two sides of the parental $S$ cell and it neighboring $S$ cells. The rearrangement of the limbal cells follows the same steps as in 1c, with a path that confined to the limbus.

ii. $P_L$ cells divide to produce a corneal $P$ cell, $Pc$. the $P_L$ offspring remains in the location of the parental $P_L$ cell in the limbus and the $P_C$ offspring becomes a corneal cell. The removed cell will be from the cornea and the location of the removed cell and the rearrangement of cells location is the same as in 1.

**Initial conditions**: The simulation is initialized as follows:

1. A square lattice grid is created with a size of ($2R$+3 X $2R$+3) while $R$ equals the radius of the cornea. A circle representing the cornea is also created around the center of the grid with the radius $R$. We used $R$=100 cells to match the *in vivo* data.
2. A 1 cell wide ring in the circumference is marked in as the limbus.
3. Each $S$ cell in the limbus is given a distinctive lineage marker, numbered from 1 to 642 (~$2\pi R$).
4. The RLS of $P$ cells is initialized to the steady-state distribution determined by running the simulation for long periods of time.
5. For visualizing the data each $S$ cell is also color-coded randomly with a color index from 1 to 5 to create the supplementary movies.

#### II. Unmixing parameter

To quantify the clone unmixing, we defined a parameter, $\phi$, that captures the deviation from a distribution where all the clones are perfect circulars sectors with a base in the limbus and an apex in the center of the cornea (Fig. S1). Consider a ring of cells (Fig. S1), In this ring, there could be several clones. The maximal angle between cells of a specific same clone $c$ is denoted by $\theta_c$, such that $0 \leq \theta_c \leq \pi$ (Fig. S1). This angle defines a ring sector $\xi_c$ The unmixing parameter of the clone $c$ in the $i^{th}$ ring is defined as,

(S1)
$$\phi_{c,i} = \frac{\text{Number of cells of clone } c \text{ in ring sector } \xi_c}{\text{Total number of cells in ring sector } \xi_c}\ .$$

If all the cells in the sector $\xi_c$ are of from clone $c$, then, $\phi_{c,i} = 1$. If the sector $\xi_c$ contains different clones, other than $c$, than $\phi_{c,i}$ is smaller than one. $\phi_{c,i}$ is always positive and approaches zero and the number of cells from the clone $c$ is small, relative to other clones in the sector. The unmixing parameter of the $i^{th}$ ring is,

(S2)
$$\phi_i = \left(\frac{1}{N_{c,i}}\sum_{c=1}^{N_{c,i}}\phi_{c,i}\right)\left(\frac{N_{c,i}}{N_{c,\text{limbus}}}\right),$$

where $N_{c,i}$ is the number of clones in the $i^{\text{th}}$ ring, and $N_{c,limbus}$ is the number of clones in the limbus. The first term in equation S2 is simply the average of $\phi_{c,i}$ on the ring while the second term penalizes the case in which the number of clones in the ring decreases relative to the number of clones in the limbus. In the case of a perfect stripe pattern (Fig. S1A), $\phi_i = 1$. In the case of mixed 'salt and paper,' pattern $\phi_i$ approaches zero. Figure S1A and figure S1B illustrates the value of $\phi$ for different patterns.

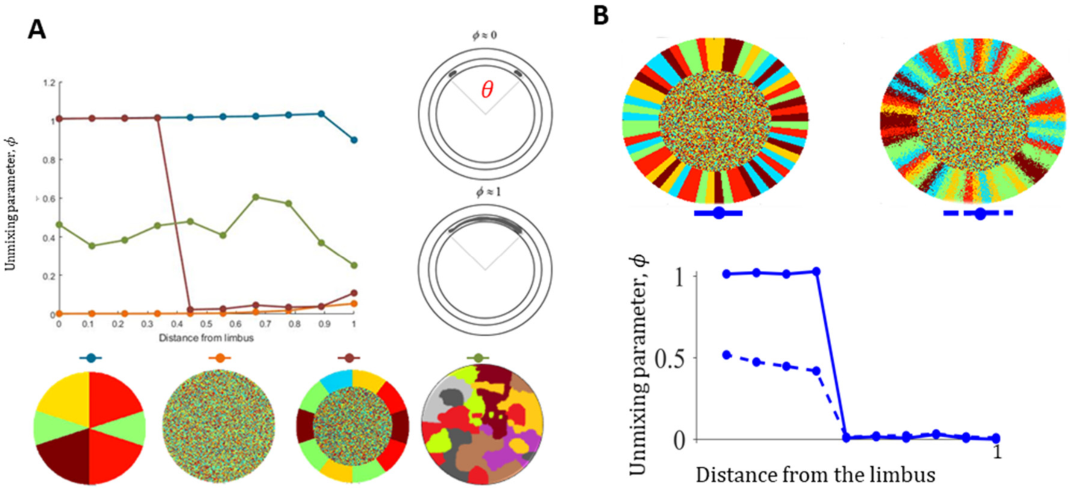


**Figure S1**: **(A)** The unmixing parameters as a function of the distance from the limbus. The right side panel demonstrates two opposing scenarios **(B)** The unmixing parameter for an ideal (left) and noisy (right) stripe pattern.

### III. Limits on coupled dynamics

**Renewal without bias.** In this case, replication and removal occur at the same local environment of a circle with a radius of *m* cells. As the cells are replicating, the lineage starts to propagate from the limbus towards the center. When $RLS = 1$, each corneal cell can replicate only once and therefore the maximal possible location of a removed cell is *m* cells from the limbus. Thus, in steady-state, after many replications, the linage front will propagate to a distance of *m* cells from the limbus.

When the RLS is increasing, the front location in steady-state will propagate further towards the center. Increasing the replicative lifespan by one will lead to a *maximal* increase of one cell in the front location (the deterministic limit). (Fig. 2B). In the deterministic limit, the front location is equal to $m + RLS$ and thus the area of the renewed cornea scales as a $C - RLS^2$, where $C$ is a constant. The actual location of the front in the simulation is indeed closer to the limbus (Fig. 2B). We captured the simulation trend by considering the propagation of the front as a stochastic process.

Consider a random variable, $x(t)$. At time interval, $\tau$,

(S3) $$x(t+\tau) = \begin{cases} x(t)+1 & p \\ x(t) & 1-p \end{cases}$$

Taking $\tau = 1$, after $N$ generation $p(x(t+N) = \chi)$ is the same as binomial distribution; having $\chi$ successes out of $N$ trials, where $p$ is the probability of success.

If this process is being carried out many times, all possible locations, $\chi$, in which $p(x(t+N) = \chi) > 0$, will be populated. That is, we define the front location in steady-state as, $\chi_s = \min(\chi \mid p(x(t+N) < \chi) = 1)$ which amounts to $\chi_s = \min(\chi \mid 1 - p(x(t+N) < \chi) = 0)$.

In the case of the tissue, the number of trials is the replicative lifespan (RLS). The new cells that are formed can replace cells in a local neighborhood with a radius of $m$ cells. When there is no centripetal bias, the probability of success depends on the geometry of the grid and the radial shape of the front. Without bias, the probability of moving forward depends on the fraction of neighbors that are not reached by the clone yet. In the case of a rectangular grid, this fraction is 3/8. Using $p = 0.35$ was used in the graph shown in Fig. 2B.

**Limit on minimal RLS for renewal.** We show in Figure 3B of the main text, that the minimal replicative lifespan required for renewal, $RLS_{min}$, decreases as a function of centripetal bias. Here we derive the theoretical lower limit on $RLS_{min}$ in the case of perfect bias.

As the front location from the limbus is smaller than $m$, the radius of the replication-removal area, the maximal propagation possible is doubling of the current distance from the limbus. And therefore the minimal number of replication it will take the front to reach a distance $m$ from the limbus is log2($m$). Once the front reached a distance $m$ from the limbus, the number of remaining replications is $RLS_{\min} - \lceil \log_2(m) \rceil$ and the maximal propagation of the front per one replication is $m$. Therefore the minimal number of replications to reach the center of the cornea is given by solving,

(S4) $$R = \lceil \log_2(m) \rceil + (RLS_{\min} - \lceil \log_2(m) \rceil) \cdot m \ ,$$

and therefore,

(S5) $$RLS_{\min} = \frac{R}{m} + \lceil \log_2(m) \rceil (1 - \frac{1}{m}) \ .$$

In the case of figure 3B, $m = 5$, $R = 100$ and therefore the limit on $RLS_{\min}$ is around 22 replications.

**Limit on minimal renewal time.** Figure 3C shows that as the centripetal bias is increased, the renewal time goes down. In the case of perfect bias, the average progress of the front is given by,

(S6) $$\langle s \rangle = \sum_{i=1}^{m} \frac{i}{m} = \frac{m+1}{2} \ .$$

Therefore, the renewal, in this case, is simply, $R / \langle s \rangle = 2R / (m+1)$.

#### IV. Limits on uncoupled dynamics

**Renewal time dependence RLS**. In figure 4C, we show the dependence of the renewal time on RLS in the case of uncoupled replication-removal. First, we assume perfect bias and *RLS* = 0. In this case, only the stem cells replicate and the cells are removed from the center of the cornea. During $t_d$, the corneal doubling time, the number of stem cells which divides asymmetrically is given by, $(\lambda_s / \lambda_p) \cdot p_a \cdot 2\pi R$. If the front is located at a distance *R* - *r* from the limbus, the average number of advances per cell in the front per one corneal doubling time is

$$(\lambda_s / \lambda_p) \cdot p_a \cdot \frac{R}{r} \ . \tag{S7}$$

Thus the time to propagate from the limbus to location *r* is given by,

$$t_0 = \sum_{r=1}^{R} \frac{1}{(\lambda_s / \lambda_p) \cdot p_a \cdot \frac{R}{r}} = \frac{\lambda_p}{\lambda_s p_a} \left( \frac{R+1}{2} \right) \ . \tag{S8}$$

In the case of $\lambda_p$=1, $\lambda_s$=0.1, $a$=0.85, $R$=100, $t_0$ is around 600.

In the case there is no bias, the removed cell location is random. Thus, the probability that the front moves towards the center each time a stem cell divides depends on whether the removed cell is within the linage area or not. Therefore, to propagate upon division is $r^2 / R^2$ and equation S8 is changed to

$$t_0 = \sum_{r=1}^{R} \frac{1}{(\lambda_s / \lambda_p) \cdot p_a \cdot \frac{R}{r} \left( \frac{r^2}{R^2} \right)} = \frac{\lambda_p}{\lambda_s p_a} \cdot R \cdot H_R \ . \tag{S9}$$

where $H_R$ is the R$^{th}$ harmonic number. In the case, for $\lambda_p$=1, $\lambda_s$=0.1, $a$=0.85 and $R$=100 $t_0$ is around 6000.

As RLS is larger than zero, corneal cell replication also contributes to the advance of the front. At low RLSs, the number of cells that replicate increases by the factor of $2^{RLS}$, and thus the renewal time scales as $2^{-RLS}$ (Fig. 3C).

**Stripe arrival time.** In the case of perfect centripetal bias, in each division, cells are removed from the center of the cornea. Therefore, the maximal distance a stripe can advance in a single replication time is doubling its length and the minimal number of divisions needed for the stripe to reach the center is $log_2(R)$ (Fig. 5B).

### V. Model parameters

| Parameter | Value and references |
|---|---|
| The radius of the cornea and basal corneal epithelial cell, *R*. | 100 Cells. (4,5) |
| Relative proliferative rates of the stem cells and progenitor cells $(\lambda_s / \lambda_p)$. | 0.1, (6,7) |
| Probability of asymmetric replication of *S* cells, $p_a$. | 0.85, Cornea (8), Epidermis (9), Esophagus (10) |
| Neighborhood radius of interacting cell in the 'coupled' model, *m*. | 5 cells (11,12) |
| The fraction of stem cells in the 'hierarchical' model, $f_s$ | 0.1 (7) |

**Table S1**: Model parameters.
